# Darker eggs resist more to desiccation: the case of melanin in *Aedes*, *Anopheles* and *Culex* mosquito vectors

**DOI:** 10.1101/109223

**Authors:** Luana C Farnesil, Helena C M Vargas, Denise Valle, Gustavo L Rezende

**Affiliations:** Laboratório de Biologia Molecular de Insetos, Instituto Oswaldo Cruz, Fiocruz, Rio de Janeiro, RJ, 21045-900, Brazil.; Laboratório de Química e Função de Proteínas e Peptídeos, Centro de Biociências e Biotecnologia, Universidade Estadual do Norte Fluminense Darcy Ribeiro, Campos dos Goytacazes, RJ, 28013-602, Brazil.; Laboratório de Biologia Molecular de Flavivírus, Instituto Oswaldo Cruz, Fiocruz, Rio de Janeiro, RJ, 21045-900, Brazil.; Instituto Nacional de Ciência e Tecnologia em Entomologia Molecular, Rio de Janeiro, RJ, 21941-902, Brazil.

**Author notes:** **Summary statement:** The ability of mosquito eggs of several species to resist differently to dry conditions is investigated. In particular, it unravels why *Aedes aegypti* eggs survive for several months outside water.

**Keywords:** *Aedes*, *Anopheles*, *Culex*, desiccation resistance, egg, embryogenesis, insect, melanin, mosquito, viability

## Abstract

Mosquito vectors lay their eggs in the aquatic milieu. During early embryogenesis water passes freely through the transparent eggshell, composed of exochorion and endochorion. Within two hours the endochorion darkens via melanization but even so eggs shrink and perish if removed from moisture. However, during mid-embryogenesis, cells of the extraembryonic serosa secretes the serosal cuticle, localized right below the endochorion, which greatly reduces water flow and allows the egg to survive outside the water. The degree of egg resistance to desiccation (ERD) at late embryogenesis varies among different species: *Aedes aegypti, Anopheles aquasalis* and *Culex quinquefasciatus* eggs can survive in a dry environment for ≥ 72, 24 and 5 hours, respectively. In some adult insects, darker-body individuals show greater resistance to desiccation than lighter ones. We asked if melanization enhances serosal cuticle-dependent ERD. Species with higher ERD at late embryogenesis exhibit more melanized eggshells. The melanization-ERD hypothesis was confirmed employing two *Anopheles quadrimaculatus* strains, the wild type and the mutant GORO, with a dark-brown and a golden eggshell, respectively. In all cases, serosal cuticle formation is fundamental for the establishment of an efficient ERD but egg viability outside the water is much higher in mosquitoes with darker eggshells than in those with lighter ones. The finding that pigmentation influences egg water balance is relevant to understand the evolutionary history of insect coloration. Since eggshell and adult cuticle pigmentation ensure insect survivorship in some cases, they should be considered regarding species fitness and novel approaches for vector or pest insects control.

## Background

Mosquitoes of the genera *Aedes, Anopheles* and *Culex* transmit pathogens that are the causative agents of diverse diseases such as dengue, chikungunya, Zika and West Nile viruses, malaria and lymphatic filariasis (Christophers, 1960; Clements, 1992; Kramer et al., 2008; Bhatt et al., 2013; Simonsen and Mwakitalu, 2013; Vega-Rua et al., 2014; Freitas et al., 2016). Blocking mosquito life cycle is an effective way to hamper disease transmission (WHO, 2013)

Mosquitoes lay their eggs in water pools, some of which are temporary (Clements, 1992). Water passes freely through their eggshells during early embryogenesis and drying of these water collections leads to egg desiccation, preventing its development. At this stage mosquito eggshell is composed of a brittle exochorion and a smooth transparent endochorion (Clements, 1992; Monnerat et al., 1999). The endochorion darkens less than three hours after being laid (Christophers, 1960; Clements, 1992), (Figure 1A) due to the melanization process.

**Figure 1:**
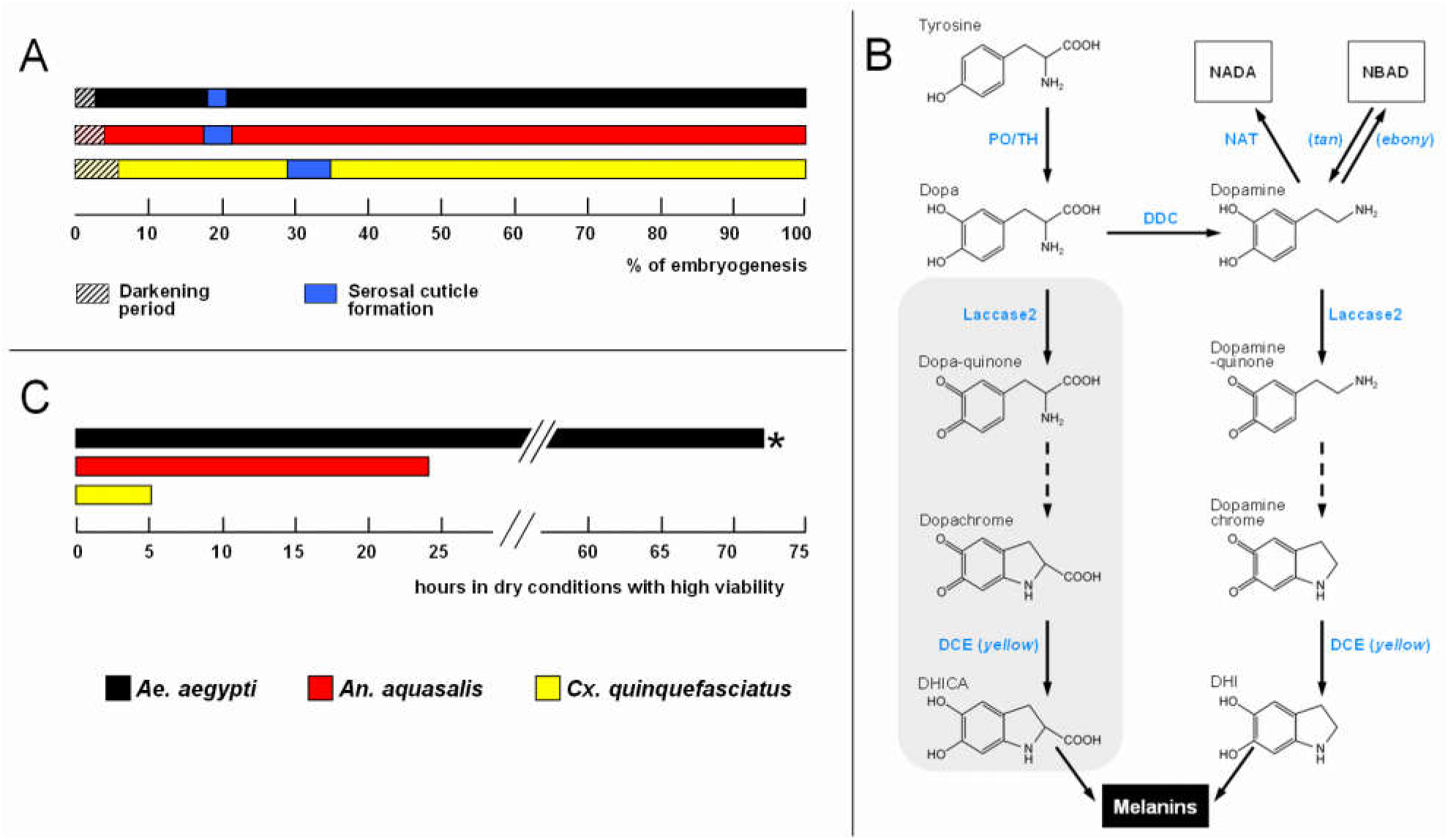
Events related to mosquito embryogenesis. (**A**) Periods of egg darkening and serosal cuticle formation. Shown as a percentage of the total embryonic development for each species, which is 77.4, 51.3 and 34.2 hours after egg laying for *Ae. aegypti*, *An. aquasalis* and *Cx. quinquefasciatus*, respectively. (**B**) Melanization pathway. Chromes are formed non-enzymatically. DHICA: 5,6-dihydroxyindole-2-carboxylic acid, DHI: 5,6-dihydroxyindole. NADA (N-acetyldopamine) and NBAD (N-β-alanyldopamine) are also substrates for Laccase 2, originating quinones that participate in the sclerotization pathway. Grey background: Dopa contribution for melanin formation is minor since (see main text). Enzyme names are shown in blue and *Drosophila melanogaster* mutants are shown in italic. PO: phenoloxidase, TH: tyrosine hydroxylase, DCE: dopachrome conversion enzyme, DDC: dopa decarboxylase, NAT: N-acetyltransferase, *tan*: N-β-alanyldopamine hydrolase, *ebony*: N-β-alanyldopamine synthase. (**C**) Egg resistance to desiccation at the end of embryogenesis. At 80% of total embryogenesis, eggs were transferred from water to dry conditions (20-55% relative humidity), and their viability monitored at regular intervals. \**Ae. aegypti* eggs are viable outside water for even longer periods (Christophers, 1960; Kliewer, 1961; Rezende et al., 2008). All data in **A** and **C** were recovered from Vargas et al. (2014), except darkening period obtained from Christophers (Christophers, 1960) and Clements (Clements, 1992).

Melanization commences with L-tyrosine hydroxylation driven by phenoloxidase or tyrosine hydroxylase that originates dopa who is decarboxylated via Dopa Decarboxylase forming dopamine. Laccase 2 act on dopa or dopamine oxidizing them and forming quinones that are further cyclized non-enzymatically giving rise to dopachrome and dopaminechrome. These two molecules are substrates for Dopachrome conversion enzyme originating DHICA and DHI that are further employed in the synthesis of the polymeric melanin. Since dopa is an inadequate substrate for Laccase2 its contribution for melanin formation is minor. Dopamine can also be β-alanylated or acetylated, originating NBAD and NADA that are further transformed in quinones that participates in sclerotization (Figure 1B) (Schlaeger and Fuchs, 1974; Li and Christensen, 1993; Johnson et al., 2001; Wu et al., 2013; Arakane et al., 2016; Rezende et al., 2016).

However, even melanized *Aedes* eggs shrink and die in a few hours if removed from moist (Rezende et al., 2008; Rezende et al., 2016). On the other hand, between 17 and 35 percent of embryogenesis occurs the production of the serosal cuticle (Figure 1A), an extracellular matrix secreted by the serosa, an extraembryonic membrane. The serosal cuticle is located below the endochorion and its formation considerably reduces water passage through the eggshell, prompting eggs to maintain their viability outside the water (Rezende et al., 2008; Goltsev et al., 2009).

Curiously, the level of egg resistance to desiccation (ERD) varies among mosquito species at the end of embryogenesis: while *Ae. aegypti* eggs can survive for at least 72 hours in a dry environment (high ERD), those of *An. aquasalis* and *Cx. quinquefasciatus* in the same condition can survive, respectively, for 24 hours (medium ERD) and 5 hours (low ERD) (Figure 1C) (Vargas et al., 2014). Physical and biochemical features of these eggs were investigated in order to identify traits related with these differences. Chitin content is directly related to ERD levels while both egg volume increase during embryogenesis and eggshell superficial density are inversely related to. Moreover, other yet unidentified traits might also be relevant (Farnesi et al., 2015).

Although the melanization increases desiccation resistance of adult insects of different orders (Kalmus, 1941; Parkash et al., 2009; Wittkopp and Beldade, 2009; King and Sinclair, 2015) it is currently unknown if the same process occurs in insect eggs. We investigated here if the intensity of eggshell pigmentation is related to desiccation resistance phenomenon in mosquito vector eggs.

## Methods

### 1. Mosquito sources and rearing

Experiments were conducted with *Aedes aegypti* (Linnaeus, 1762), *Anopheles aquasalis* (Curry, 1932) and *Culex quinquefasciatus* (Say, 1823) continuously maintained at the Laboratório de Fisiologia e Controle de Artrópodes Vetores (LAFICAVE), Instituto Oswaldo Cruz, Rio de Janeiro, RJ, Brazil and the strains ORLANDO and GORO of *Anopheles quadrimaculatus* (Say, 1824), reared between March and August 2013 at the Florida Medical Entomology Laboratory (FMEL), Florida University, Vero Beach, FL, USA. Both *An. quadrimaculatus* strains, ORLANDO (MRA-139) (https://www.beiresources.org/Catalog/BEIVectors/MRA-139.aspx - acessed 15 February 2016) and GORO (MRA-891) (https://www.beiresources.org/Catalog/BEIVectors/MRA-891.aspx - acessed 15 February 2017) were obtained through the Malaria Research and Reference Reagent Resource Center (MR4) (Manassas, VA, USA), as part of the BEI Resources Repository, NIAID, NIH and were deposited by MQ Benedict. The *An. quadrimaculatus* ORLANDO strain is mentioned in this work as "WT" (i.e. wild type). The *An. quadrimaculatus* GORO strain contains two EMS-induced mutations, both on the X chromosome, and was generated crossing the GOCUT strain (MRA-123) and the ROSEYE strain (MRA-122). GORO genotype is go^1 pk^ + ro^1 and its phenotype, as seen in Figure 3A-D, is golden cuticle at all stages and rose eye from larvae on. Larvae were reared at 26 ± 1 °C in rectangular plastic basins (*Ae. aegypti, An. aquasalis* and *Cx. quinquefasciatus*) or rectangular iron pans coated with vitreous enamel (*Anopheles quadrimaculatus*) containing 300 specimens within 1 liter of water and with 1 gram of food being provided every two days. Water and diet source varied in each case: dechlorinated water and cat food Friskies^®^ (“Peixes – Sensações marinhas”, Purina, Camaquã, RS, Brazil) for *Ae. aegypti* and *Cx. quinquefasciatus*, brackish dechlorinated water (2 mg of marine salt/mL of dechlorinated water) and fish food Tetramin^®^ (*Tetramarine Saltwater Granules, Tetra GmbH, Germany*) for *An. aquasalis*, tap water and brewer’s yeast/liver powder (1:1) for *An. quadrimaculatus*. In all cases, adults were kept at 26 ± 1 °C, 12/12 h light/dark cycle, 70 - 80% relative humidity and fed *ad libitum* with 10% sucrose solution.

### 2. Synchronous egg laying

The synchronous egg laying method was adapted from Valencia *et al*. (Valencia et al., 1996b; Valencia et al., 1996a), as previously described (Rezende et al., 2008; Vargas et al., 2014; Farnesi et al., 2015). For egg production, females of all species, three to seven days old, were sugar deprived for 24 hours and then blood-fed on anaesthetized chickens (*An. quadrimaculatus*) or guinea pigs (all other species). Gravid mosquito females were transferred to 15 mL centrifuge tubes and anesthetized in ice for a few minutes. The interval between blood meal and egg laying induction, as well as the procedure adopted for obtaining eggs, varied according to the species.

*Aedes aegypti* and all anopheline females were anaesthetized in ice three to four days after blood feeding. Groups of five to ten sleeping females were then rapidly transferred to upside down 8.5 cm diameter Petri dishes, where the lid became the base. This base was internally covered with Whatman No. 1 filter paper. After the females were awaken, a process that took 3-10 minutes, the filter paper was soaked with the same water employed to rearing each species, thus stimulating the laying of the eggs that were deposited individually or in small disorganized groups.

Groups of five to ten *Culex quinquefasciatus* females were anaesthetized in ice five to six days after the blood meal and then transferred to 8.5 cm diameter Petri dishes in the normal position (not upside down) without filter paper. After insect recovery, dechlorinated water was added with the aid of a micropipette through a small hole in the lid until the females were pressed against it, which stimulated egg laying. A second small hole was present in the lid to allow air outlet while water was being introduced. Eggs were deposited in organized rafts containing from few dozens to hundreds of eggs.

In all cases egg laying lasted one hour in the dark, inside an incubator at 25 ± 1 °C. Petri dishes were then opened inside a large cage where the females were released. Eggs were allowed to develop at 25 °C until being employed in the experiments. For *Ae. aegypti* and anopheline eggs the sides of the Petri dishes were sealed with parafilm, in order to avoid water evaporation. For *Cx. quinquefaciatus* eggs, rafts were kept intact prior to the first experimental point, when they were transferred to Petri dishes whose base was covered with Whatman No. 1 filter paper soaked with dechlorinated water. Rafts were carefully disrupted and the eggs were spread with the aid of a painting brush.

The procedure and use of live chicken followed the UF-IACUC Protocol no. 201003892. The procedure and use of anaesthetized guinea pigs was reviewed and approved by the Fiocruz institutional committee ‘Comissão de Ética no Estudo de Animais’ (CEUA/FIOCRUZ), license number: L-011/09.

### 3. Eggshell darkening analysis in *Ae. aegypti, An. aquasalis* and *Cx. quinquefasciatus*

Eggs at approximately 80% of embryogenesis completion had their exochorion removed with bleach (NaOCl, 6% active chlorine) treatment for one minute followed by three washes with dechlorinated water. These exochorion-depleted eggs were then kept in moist filter paper until hatching. Eggshells were then transferred into a microscopy slide and brightfield images were obtained with a digital imaging acquisition system coupled to a Zeiss Axio Scop 40 microscope. Two experiments per species were performed, each one consisting of at least 9 eggshells. The image acquisition setup was the same, in both the microscope and the computer, for all images. Eggshell melanization degree was evaluated employing the ImageJ software (https://imagej.nih.gov/ij/) with the 'Measure' function within the 'Analyze' menu. This function calculates the mean densitometric value of the selected area in a 8-bit grey scale, i.e. a completely white and a completely black pixel has, respectively a value of 255 and 0. Representative circular regions were selected, always close to the hatching line (see lined circles on Figure 2). The densitometry of each eggshell was subtracted against the densitometry of a fixed circular region of non-saturated white background (with a value of 232). Densitometry values were then inversed (i.e. a white and a black pixel measuring, respectively, 0 and 255) and darkening percentages were calculated, assuming the mean value of *Ae. aegypti* eggshells as 100%.

**Figure 2:**
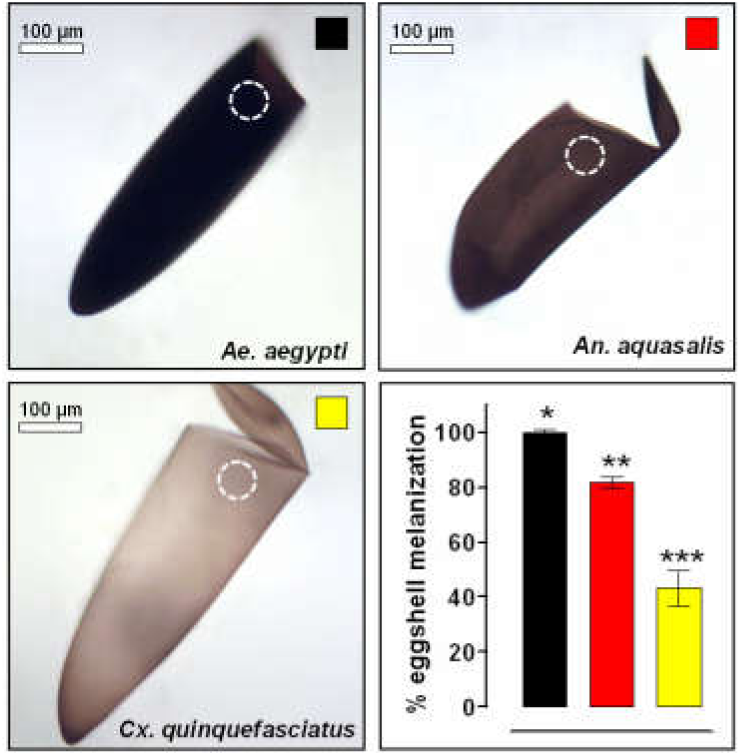
Mosquito eggshell melanization varies among species. Melanization degree was quantified in empty eggshell images obtained with bright field microscopy employing the ImageJ software (lower right graphic). The maximum melanization level was arbitrarily attributed to *Ae. aegypti* eggshells. The measured region, always near the hatching line, is indicated by white circles. A direct correlation between melanization and ERD degree occurs (compare with Figure 1). Values represents the mean ± s.d. of two experiments, each consisting of at least 9 eggshells. All observed differences are statistically significant (Kruskal-

### 4. Detection of serosal cuticle formation in *An. quadrimaculatus*

Serosal cuticle synthesis was evaluated in both WT and GORO strains of *An. quadrimaculatus* employing two approaches: air drying and bleach treatment, as previously described for the other mosquitoes (Rezende et al., 2008; Goltsev et al., 2009; Vargas et al., 2014).

For the air drying assay, replicates consisting of 30 synchronized eggs at distinct stages of embryogenesis (comprising seven time points in total, see *x*-axis in the Figure 3E) were blotted onto a dry Whatman No. 1 filter paper to remove all water. Eggs were then left drying on air for 15 minutes, when shrunken or intact eggs were counted under a stereomicroscope. For each time point, three independent experiments were performed, each with 30 eggs, for each strain. Experiments were performed at 25 ºC and the relative humidity varied between 65 and 75%.

**Figure 3:**
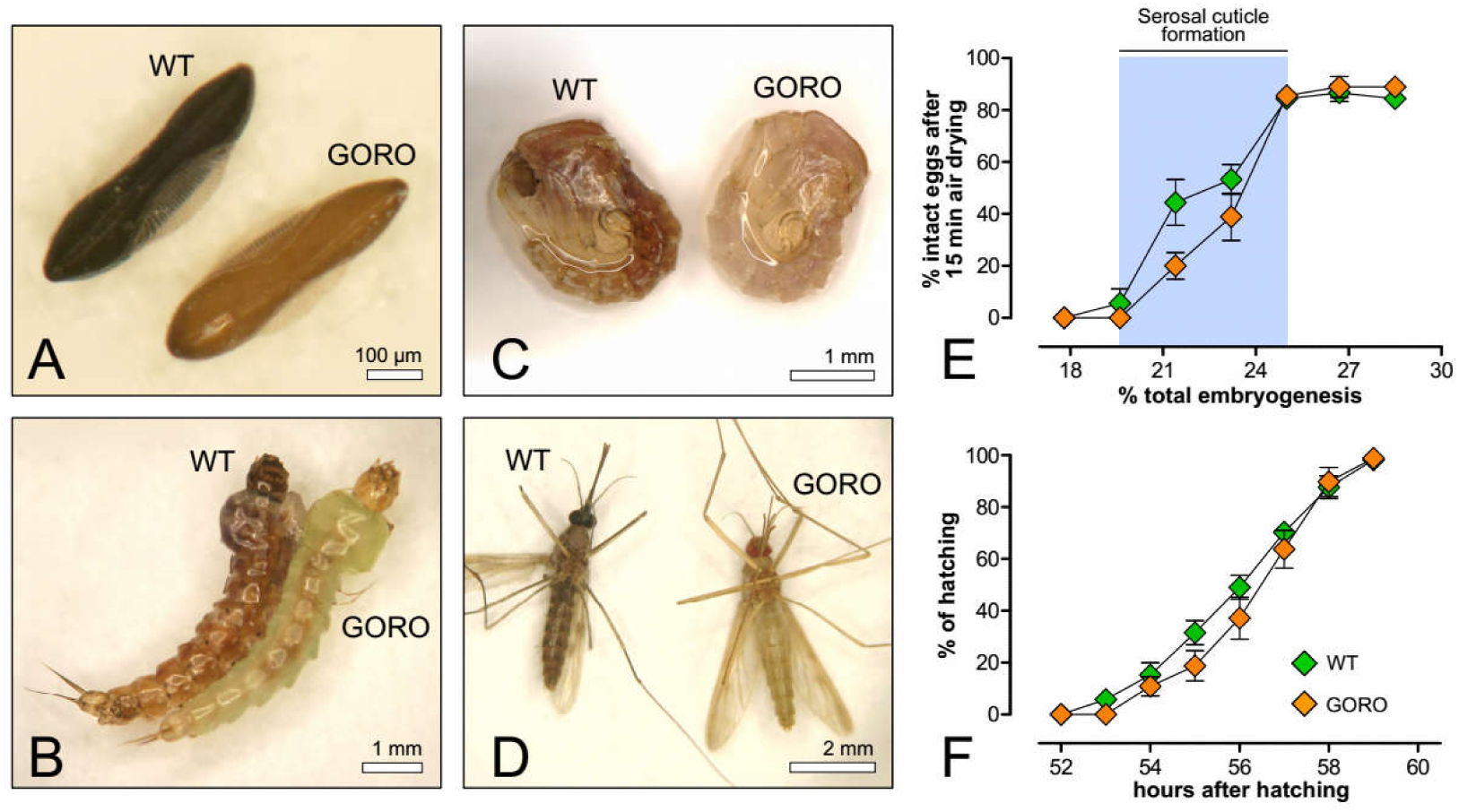
Embryogenesis of the weakly pigmented *Anopheles quadrimaculatus* GORO strain proceeds similarly to the WT. GORO means ‘GOlden cuticle and ROse eyes’. (**A**) eggs, (**B**) larvae, (**C**) pupae and (**D**) adults. (**E**) Eggs at different embryonic ages developing at 25 ºC were airdried for 15 minutes and the percentage of eggs that did not shrink (i.e. intact eggs) was then registered. Relative humidity ranged between 65 and 75%. The abrupt alteration in egg permeability, highlighted by a blue stripe, is coupled with serosal cuticle formation (see Figure 4). Points represent mean ± s.e. of three independent experiments, each one with 30 eggs per point (total of 630 eggs per strain) (**F**) Cumulative larval hatching at 25 °C; data were normalized by total eclosion, obtained 24 hours after the expected embryogenesis completion. Each curve represents mean and standard error of three independent experiments consisting of 120 eggs each (total of 360 eggs per strain).

Prolonged incubation with bleach digests both the egg exochorion and endochorion while leaving the serosal cuticle intact. Synchronized *Anopheles quadrimaculatus* eggs from both strains were treated with bleach (6% active chlorine) during 3 - 10 min at different stages of embryogenesis, before and after the abrupt change in egg permeability (detected through the air drying experiment described above). The resulting material was analyzed under a stereomicroscope (MIA 3XS S/N 0342, Martin Microscope Company) with an Olympus U-CMAD3 U-TV1X 2 adapter and Nikon CodPix 5400 camera, coupled with a digital image acquisition system. For each strain and time point two independent experiments, each with at least 20 eggs, were performed.

### 5. Definition of the end point of *An. quadrimaculatus* embryogenesis

The total period necessary for embryonic completion in both WT and GORO strains was defined as previously described for other mosquitoes (Farnesi et al., 2009; Vargas et al., 2014). Two hours before the (empirically) estimated hatching of the putative first larva, eggs were flooded with a solution of 150 mg/ 100 mL yeast extract (SIGMA # Y1625) prepared in tap water. Egg eclosion was counted hourly, until no more hatchlings were observed. Twenty four hours after the eclosion of the last putative larvae the samples were checked again to confirm that total hatching was recorded. The embryogenesis end point was defined as the period necessary to hatch 50% of total larvae. For each strains, three independent experiments, each with 120 eggs, were performed.

### 6. Embryo viability under dry conditions

All species and strains were employed in this experiment. In each case, groups of 40 or 50 synchronized eggs, obtained as explained above (section 2 of Methods), were removed from water and blotted onto dry Whatman Nº 1 filter paper with the aid of a paint brush, at specific moments of embryogenesis (see Figure 4 for details). Eggs remained developing in this dry environment for 2, 5, or 10 hours. After these periods, eggs were transferred back to moist conditions until embryogenesis completion. In all experiments the total test interval ("wet-dry-wet") was shorter than the period necessary for embryogenesis completion ((Vargas et al., 2014); Figure 3F). Egg viability was quantified through larval hatching, induced with 150 mg/ 100 mL yeast extract solution (Farnesi et al., 2009; Vargas et al., 2014), prepared with the same water used for rearing (section 1 of Methods). Larval eclosion was recorded hourly until no more hatchlings were observed for two successive hours. Total larval hatching was confirmed 24 hours later.

**Figure 4:**
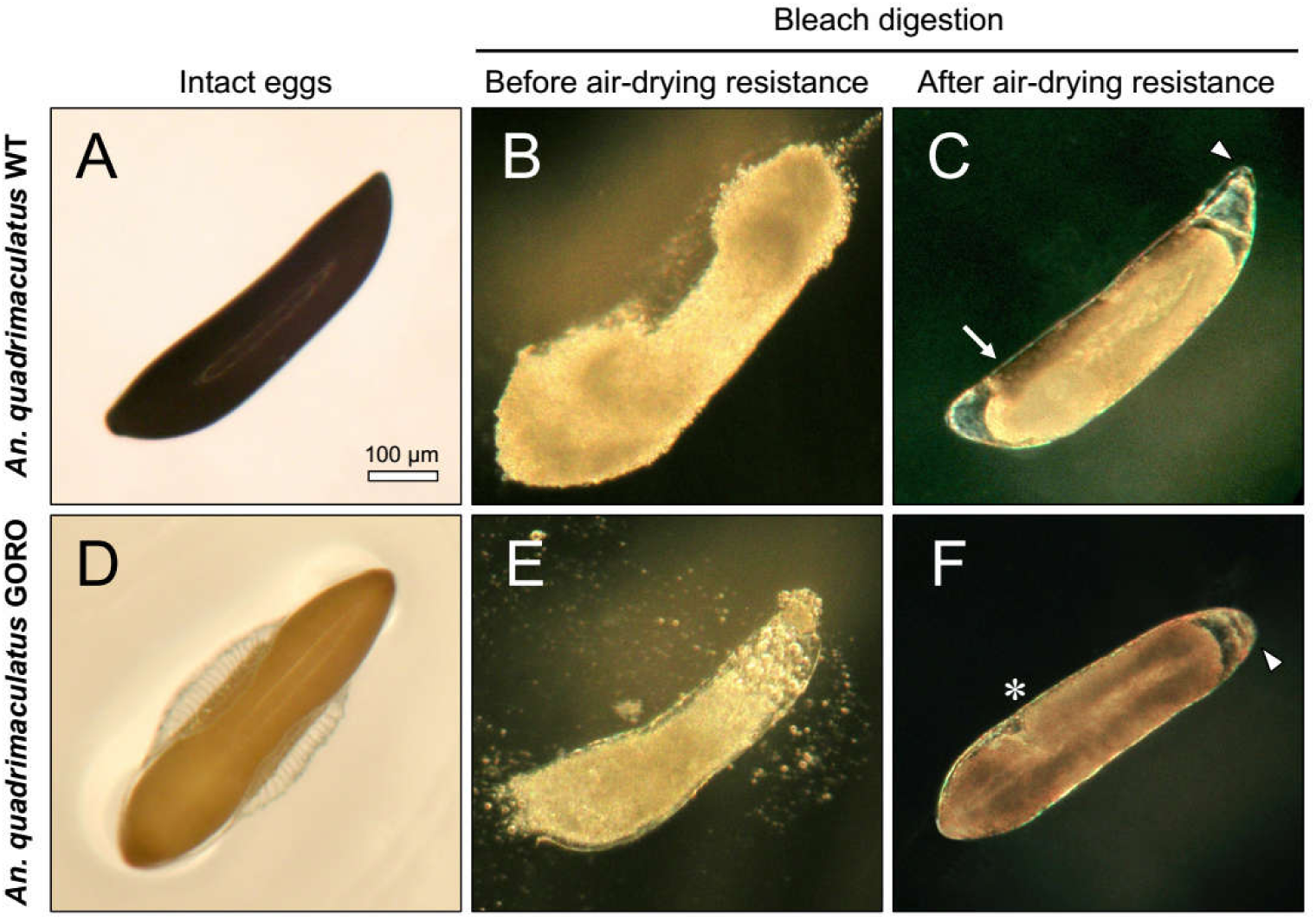
Resistance to air-drying is related to serosal cuticle formation in both *An. quadrimaculatus* strains. Serosal cuticle presence was determined by chorion digestion driven by bleach (6% active chlorine). (**A, D**) Intact eggs. (**B, E**) Eggs treated with bleach before acquisition of air-drying resistance are totally digested while (**C, F**) eggs exposed to the same procedure after acquisition of air-drying resistance remain intact due to the presence of the serosal cuticle (see Figure 3E). Arrow: endochorion remnants not yet digested; arrowheads: serosal cuticle boundaries; asterisk: posteriormost end of the germ band. All images are in the same magnification.

Viability control samples containing at least 120 eggs, kept continuously in moist filter paper until the end of embryogenesis, were employed in all cases. Experimental data were normalized with these controls, whose hatching was induced with yeast extract solution (150 mg/ 100 mL).

Three independent experiments were performed for each species or strain, using triplicates at least, inside an incubator at 25±1 °C. Relative humidity varied between 60 and 80% for both *An. quadrimaculatus* strains and between 20 and 55% for all other species.

### 7. Statistical analysis

For the analysis of eggshell darkening, air drying, embryogenesis period and embryonic viability under dry conditions the adequate sample size (n) of each experiment was defined from preliminary experiments. For all these experiments, eggshells or eggs were randomly collected from the filter paper (see item 2 of Methods). Outliers were removed after Dixon’s Q test. Kruskal-Wallis Nonparametric Test (P< 0.0001) was used in eggshell melanization analysis, One Way Analysis of Variance (ANOVA) followed by Tukey’s Multiple Comparison Test (P< 0.05) was used in the egg viability experiments and the Student’s t-test (P <0.001) was used to compare viability between the two *Anopheles quadrimaculatus* strains. All statistical analyzes, except Dixon's Q test, were made using GraphPad Prism version 5.00 for Windows (GraphPad Software, San Diego, California USA, www.graphpad.com).

## Results

### Levels of eggshell melanization and egg resistance to desiccation (ERD) are directly related among species

The ERD, defined as the capacity of an egg to sustain its viability outside the water (Hadley, 1994; Gibbs et al., 1997), varies among mosquito species at the end of embryogenesis (Figure 1) (Vargas et al., 2014). In order to evaluate if these viability differences could be explained by egg pigmentation, the degrees of melanization of hatched eggshells of *Ae. aegypti*, *An. aquasalis* and *Cx. quinquefasciatus* were assessed (Figure 2). Eggs of *Aedes aegypti* and *An. aquasalis* present a homogeneous pigmentation, while *Cx. quinquefasciatus* eggs are more pigmented near its extremes. In spite of this, overall, *Aedes aegypti* exhibits the greater eggshell pigmentation, followed by *An. aquasalis* and *Cx. quinquefasciatus*.

Although establishing a direct relationship between eggshell pigmentation and ERD is tempting, other eggshell related factors, such as differences in thickness or components of the endochorion or the serosal cuticle, might account for this distinctness (Harwood, 1959; Christophers, 1960; Clements, 1992; Monnerat et al., 1999; Farnesi et al., 2015). Moreover, since we are studying mosquitoes of different genus, whose common ancestor occurred ~ 217 million years ago (Reidenbach et al., 2009), embryological and egg traits vary considerably (Vargas et al., 2014; Farnesi et al., 2015) and may not be comparable. In order to directly evaluate the relationship between melanization and ERD without any other confounding factor, we took advantage of a mutant strain of the species *Anopheles quadrimaculatus*, which shows a significant melanization deficit: the GORO strain.

### *An. quadrimaculatus* GORO embryogenesis is normal, despite its impaired melanization

The mosquito *Anopheles quadrimaculatus* is endemic to the Eastern part of North America, being a primary vector of malaria in this region (Reinert et al., 1997). The wild type strain of this species presents a dark-brown, melanized eggshell and a dark-brown cuticle in larval, pupal and adult stages (Figure 3A-D). On the other hand, the GORO strain carries a *golden cuticle* mutation within a *rose eye* background (hence the name GORO: GOlden cuticle + ROse eyes), which causes poor body melanization in all life stages (Mazur et al., 2001; BEI, 2016a; BEI, 2016b), see Methods, (Figure 3A-D). In order to assess whether the lack of proper melanization compromises embryogenesis, two embryonic traits were analyzed in WT and GORO: the chronology of serosal cuticle formation and the completion of embryogenesis (Figure 3E, F, Figure 4). Serosal cuticle formation, assessed through the abrupt acquisition of resistance to egg shrinkage (Figure 3E) and bleach digestion (Figure 4), as previously described in other mosquito species (Rezende et al., 2008; Vargas et al., 2014), occurs in between 19.6 and 25% of total embryogenesis, at the stage of complete germ band elongation (Figure 4), in both strains. Likewise, the period necessary for entire embryogenesis, approximately 56 hours after egg laying, is similar in both strains (Figure 3F). Therefore, the lack of melanization in the *An. quadrimaculatus* GORO mutant does not compromise neither serosal cuticle formation nor the total period necessary for embryogenesis completion.

### Egg resistance to desiccation after serosal cuticle formation is enhanced by melanization

The interspecific difference in egg viability when these are placed outside the water at late embryogenesis (Figure 1C) (Vargas et al., 2014) might be due to factors other than the eggshell and its serosal cuticle. For instance, it could be caused by specific metabolites inside the egg or present in the pharate larvae, such as glycerol, trehalose, glycogen or triacylglycerols, or to significant variation in the larval cuticle structure (Hinton, 1981; Hadley, 1994; Gibbs et al., 1997; Sawabe and Mogi, 1999; Gray and Bradley, 2005). Thus, we uncoupled serosal cuticle participation in ERD from other factors. Eggs from the different mosquito species and strains were removed from the water at different stages of early embryogenesis and left developing outside the water for two, five or ten hours. Hatching rates were assessed at the end of embryogenesis (Figure 5 and Table 1). In all cases serosal cuticle formation significantly increases egg viability outside the water (ANOVA followed by Tukey’s Multiple Comparison Test, P < 0.05). The role of the serosal cuticle on ERD of *Ae. aegypti* left up to ten hours in a dry environment is partial: the serosal cuticle elevates embryo viability from 30-50% before its formation to 68-81% right after its synthesis. However, all *Ae. aegypti* eggs die if remaining outside the water for 25 hours prior to serosal cuticle formation (Rezende et al., 2008). In *Anopheles* species and strains the serosal cuticle formation is essential: egg viability in dry conditions is null before, but increases considerably after serosal cuticle synthesis, as previously described for *An. quadrimaculatus* (Darrow, 1949) and *An. gambiae* (Goltsev et al., 2009). In both *Ae. aegypti* and *An. aquasalis* the hatching rate in each stage is equivalent for all dry exposure periods. Regarding *Cx. quinquefasciatus*, 20% of the eggs left outside the water for two hours before serosal cuticle synthesis survive but similar aged eggs exposed to a dry environment for longer periods do not resist. Moreover, egg viability after serosal cuticle formation is inversely proportional to the exposure period outside the water. Interestingly, in both *Cx. quinquefasciatus* and *Ae. aegypti*, a gradual increase in embryo viability was observed after serosal cuticle formation, suggesting this structure follows a process of maturation until it becomes completely functional. Regarding *An. quadrimaculatus*, in both strains the percentage of viable eggs is inversely related to the dryness period. In all conditions after serosal cuticle formation, GORO eggs are far more sensitive to dehydration than wild type ones (Student’s t-test, P < 0.001). For instance, at 25% of total embryogenesis and when left for 5 hours in a dry environment, the hatching rate of WT and GORO strains are, respectively, 85 and 17%.

**Figure 5:**
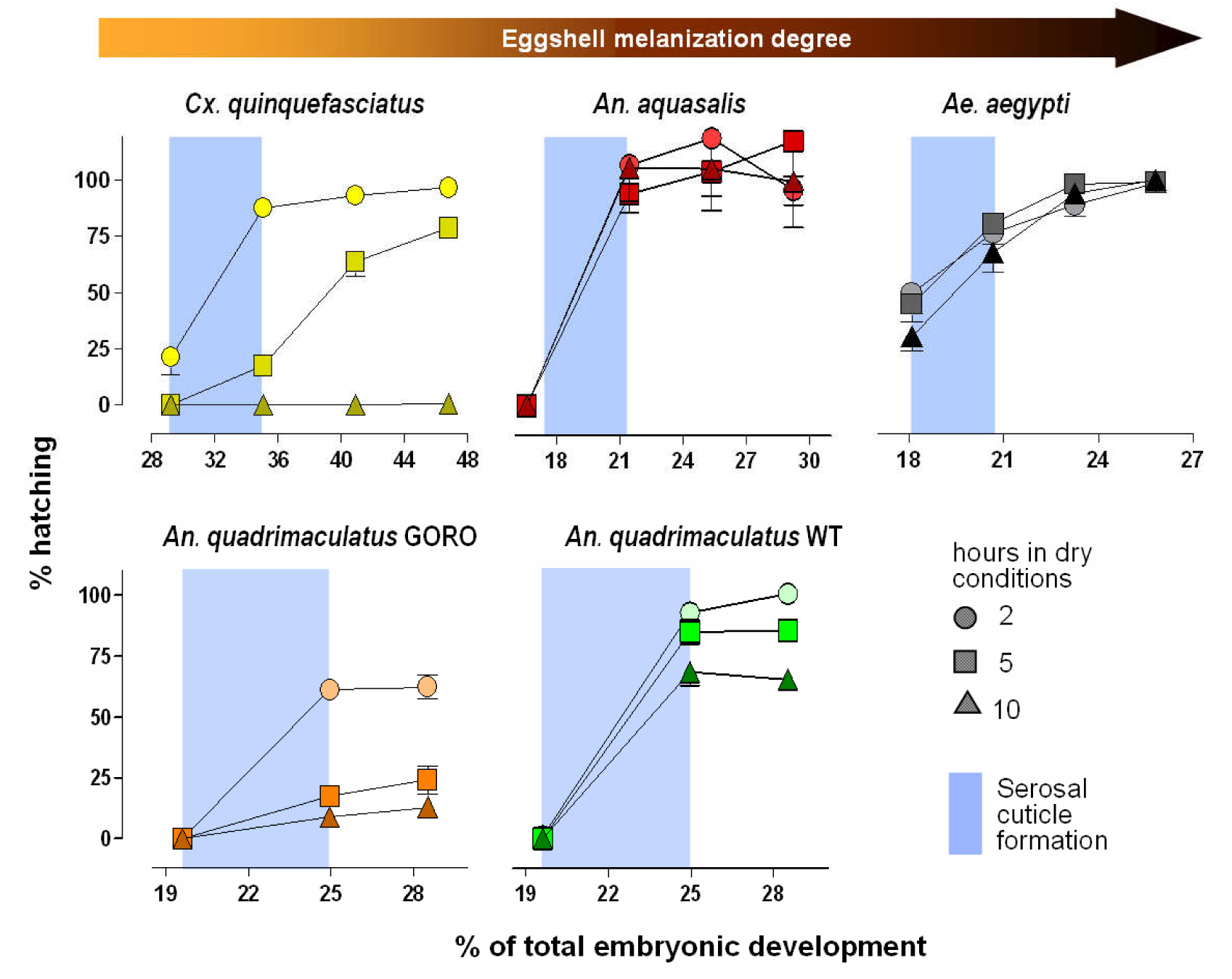
Mosquitoes with darker eggshells resist more to desiccation. Mosquito eggs were laid on water. Values in the *x*-axis indicate the moment that eggs were transferred to dry conditions, staying outside the water for 2, 5 or 10 hours. Eggs were then returned to moist filter paper until completion of embryo development, when hatching rates were evaluated. Data were normalized regarding to control samples, kept on moist conditions throughout development. Blue stripes indicate the serosal cuticle formation period (Figures 1 and 3). Each point represents mean ± s.e. of three independent experiments consisting of at least 120 eggs each. A total of at least 3240 eggs were employed for each species or strain. In all cases viability was significantly different between the two first experimental points (i.e. before and after serosal cuticle formation) (ANOVA followed by Tukey’s test, P < 0.05, see Table 1); the exception being *Cx. quinquefasciatus* at 10 hours in dry conditions. After serosal cuticle formation, *An. quadrimaculatus* GORO eggs were less viable than WT ones in all equivalent conditions (Student’s t-test, P < 0.001). All experiments were conducted at 25 ºC and relative humidity of 60-80% (*An. quadrimaculatus*) or 20-55% (other species).

**Table 1:** Egg viability of mosquito species and strains under dry conditions during embryogenesis, before and after serosal cuticle (SC) formation.

| Species or strain | Hours under dry conditions | Stage of embryogenesis<sup>#</sup> |  |  |  |
| --- | --- | --- | --- | --- | --- |
|  |  | Before SC formation | After SC formation I | After SC formation II | After SC formation III |
| *Ae. aegypti* | 2 | 50.0 ± 32.4<sup>a</sup> | 76.4 ± 16.9<sup>b</sup> | 88.9 ± 16.8<sup>b</sup> | 98.9 ± 11.5<sup>b</sup> |
|  | 5 | 44.9 ± 28.3<sup>a</sup> | 80.6 ± 10.5<sup>b</sup> | 98.3 ± 15.6<sup>b</sup> | 99.1 ± 14.1<sup>b</sup> |
|  | 10 | 30.2 ± 21.3<sup>a</sup> | 67.9 ± 30.4<sup>b</sup> | 94.3 ± 13.4<sup>c</sup> | 100.0 ± 17.9<sup>c</sup> |
| *An. aquasalis* | 2 | 0.0 ± 0.0<sup>a</sup> | 107.8 ± 33.4<sup>b</sup> | 119.7 ± 56.7<sup>b</sup> | 96.7 ± 11.5<sup>b</sup> |
|  | 5 | 0.0 ± 0.0<sup>a</sup> | 94.8 ± 24.9<sup>b</sup> | 104.8 ± 52.2<sup>b</sup> | 118.6 ± 47.3<sup>b</sup> |
|  | 10 | 0.0 ± 0.0<sup>a</sup> | 106.6 ± 44.6<sup>b</sup> | 106.2 ± 36.8<sup>b</sup> | 100.3 ± 31.1<sup>b</sup> |
| *Cx. quinquefasciatus* | 2 | 21.1 ± 22.8<sup>a</sup> | 87.6 ± 12.3<sup>b</sup> | 93.1 ± 10.2<sup>b</sup> | 96.7 ± 7.3<sup>b</sup> |
|  | 5 | 0.0 ± 0.0<sup>ha</sup> | 17.4 ± 8.8<sup>b</sup> | 63.7 ± 19.4<sup>c</sup> | 78.7 ± 10.3<sup>c</sup> |
|  | 10 | 0.0 ± 0.0<sup>a</sup> | 0.0 ± 0.0<sup>a</sup> | 0.0 ± 0.0<sup>a</sup> | 0.5 ± 0.9<sup>b</sup> |
| *An. quadrimaculatus* WT | 2 | 1.0 ± 2.9<sup>a</sup> | 92.9 ± 11.4<sup>b</sup> | 106.6 ± 10.2<sup>b</sup> |  |
|  | 5 | 0.0 ± 0.0<sup>a</sup> | 84.8 ± 14.9<sup>b</sup> | 85.5 ± 13.1<sup>b</sup> | N.D. |
|  | 10 | 0.0 ± 0.0<sup>a</sup> | 68.5 ± 16.9<sup>b</sup> | 65.3 ± 11.3<sup>b</sup> |  |
| *An. quadrimaculatus* GORO | 2 | 0.0 ± 0.0<sup>a</sup> | 60.9 ± 6.4<sup>b*</sup> | 62.0 ± 14.0<sup>b*</sup> |  |
|  | 5 | 0.0 ± 0.0<sup>a</sup> | 17.2 ± 9.2<sup>b*</sup> | 23.9 ± 16.8<sup>b*</sup> | N.D. |
|  | 10 | 0.0 ± 0.0<sup>a</sup> | 8.7 ± 7.4<sup>b*</sup> | 12.8 ± 7.1<sup>b*</sup> |  |

<sup>#</sup> The stages of embryogenesis are indicated in the x-axis of Figure 5.
Values represent mean and standard deviation of at least three independent experiments for each species and period under dry conditions.
Every experiment employed a total of at least 120 eggs for each point and for each species or strain.
Hatching percentages were normalized according to control samples kept moist throughout development.
Different letters represent significant differences among the distinct stages of embryogenesis in the same drying period and for the same species or strain (ANOVA, followed by Tukey's test P<0.05).
Asterisk means significant differences between *An. quadrimaculatus* WT and GORO strains in the same drying period and for the same stages of embryogenesis (Student's t-test, P <0.001).
N.D.: Not determined.

## Discussion

### Regarding the *Anopheles quadrimaculatus* GORO strain

Thanks to the existence of the *An. quadrimaculatus* GORO strain it was possible to prove that egg resistance to desiccation in mosquitoes is heavily dependent on serosal cuticle formation and, at the same time, that eggshell melanization positively impacts the egg survivorship outside the water. Although this interesting strains exists for at least 15 years (Mazur et al., 2001), this is the first report employing it and the genetics of the *golden cuticle* mutation present in the *An. quadrimaculatus* GORO is currently unknown. Given that melanization is also related with immunity, it would be interesting to evaluate how the GORO strain responds immune challenges in adults, larvae and eggs (Jacobs and van der Zee, 2013; Jacobs et al., 2014).

It is worth mentioning that it would not be possible to use the same approach, at least with *Aedes* mosquitoes: the mutants *bronze* and *gray*, presenting altered egg color, are embryonic lethal (Craig and A., 1967), as well as gene silencing for *Laccase 2* (Wu et al., 2013). The administration of α-MDH or Benserazide, drugs that inhibit Dopa decarboxylase activity, impedes eggs to darken completely, rendering tanned eggs (similar in color to GORO eggs); but these less melanized eggs are inviable (Schlaeger and Fuchs, 1974; Martins, 2002).

### The role of egg color in insects

Insect eggs occur in a myriad of colors, ranging from white to black with tones of yellow, orange, red, pink, green and brown, among others. Egg color may occur uniformly or in patches throughout the eggshell, or can appear in restricted areas (Hinton, 1981). These colors are produced by pigments such as melanins, sclerotins, ommochromes, pteridines, carotenoids and flavonoids (Wittkopp and Beldade, 2009; Nijhout, 2010).

Egg colors are associated with defense strategies against predators, such as homochromy, mimicry, camouflage, visual disruption and warning (aposematic) signaling (Hinton, 1981). Females of the bug *Podisus maculiventris* selectively control egg color during oviposition: darker and lighter eggs are laid on the upper and lower surface of leaves, respectively. The dark pigment protects eggs against the deleterious effects of UV light emitted from the sun (Abram et al., 2015).

This list is further expanded with melanin participation in the egg resistance to desiccation (ERD). The ERD trait has been associated with the staggering adaptive success insects show on land (Zeh et al., 1989; Jacobs et al., 2013). Two questions arise from the above considerations. A direct exposition to sunlight also increases evaporation of eggs (Hinton, 1981): does the dark pigment selectively present in eggs of *P. maculiventris* also protects against desiccation? In relation to the other non-melanin pigments; do they also protect insect eggs and cuticles in post-embryonic life stages from water loss?

### Melanin and desiccation resistance in adult insects

The melanin contribution for desiccation resistance has been previously described in adult insects: Kalmus (Kalmus, 1941) compared the desiccation resistance in adults of wild type and *yellow*, *ebony* and *black* mutants of the *Drosophila melanogaster* fly. The wild type cuticle is melanized, the cuticle of *yellow* mutants is light brown/yellowish (i.e. with a tanned color) and the cuticle of *black* or *ebony* mutants are darker than wild type ones. The more melanized a fly is, the more it resists desiccation. The *yellow* gene is related with the activity of Dopachrome conversion enzyme, required for proper melanin formation while both *black* and *ebony* genes code for enzymes necessary for NBAD production, driving dopamine usage for sclerotization, instead of melanization (Figure 1B) (Arakane et al., 2016). The same pattern was found in distinct species and morphs of *Hemideina* wetas from New Zealand and morphs of *D. melanogaster* from the Indian subcontinet: darker adults resist more against desiccation (Parkash et al., 2009; King and Sinclair, 2015). In the beetle *Tribolium castaneum* silencing of the gene *yellow-e* (*TcY-e*) leads to desiccation sensitivity of adults. These adults survive when reared at high humidity but, intriguingly, develop a slightly darker cuticle (Noh et al., 2015).

On the other hand, populations of *D. melanogaster* artificially selected for increased pigmentation does not resist desiccation more than control flies (Rajpurohit et al., 2016). This apparent incoherence might be due to other factors, since the reduction in the rate of water loss by the cuticle is one out of the three aspects of the desiccation resistance (see below). Other explanation could be related with the physicochemical properties of the melanin produced.

### How does melanin protects insect structures against desiccation?

Melanin might protects against desiccation due to its covalent or noncovalent interaction with other biomolecules such as proteins and chitin (Arakane et al., 2016). If this is the case, this association is distinct from sclerotization-driven crosslinking: both *black* and *ebony D. melanogaster* mutants present a cuticle that is less stiffen and puncture-resistant than wild type ones (Andersen, 2012). Similarly, the elytral cuticle of *T. castaneum black* mutants are more viscous and less stiffen than wild type ones (Arakane et al., 2009).

Another hypothesis is that melanin might be hydrophobic and thus hamper water flux through the cuticle, as recently suggested (Rajpurohit et al., 2016). Although both melanin precursors (DHICA and DHI, Figure 1B) are hydrophilic compounds, the molecular structure of melanin polymers varies depending on the biochemical conditions of polymerization and, therefore, "melanin" is a diffuse term for a rather diverse group of complex pigments (Prota, 1992; Ito et al., 2011; d'Ischia et al., 2013; Shamim et al., 2014; Arakane et al., 2016). In fact, exists in the literature descriptions of melanin being both water-soluble (Mostert et al., 2010) and water-insoluble (Shamim et al., 2014). Thus the *D. melanogaster* darker-selected populations might not have a higher desiccation resistance (Rajpurohit et al., 2016) due to the production of "hydrophilic melanins" in this specific situation.

In any case, although melanization in some instances increases desiccation resistance, as shown in the present work, this is not an universal rule (Wigglesworth, 1948), as further exemplified below for other insect eggs.

### Melanin localization in the eggshell and other egg traits related with resistance to desiccation

In any organism, an increase in resistance to desiccation is related with three aspects: a higher initial body water store, a reduction in the rate of water loss and an increase in the tolerance to water loss (Hadley, 1994; Gibbs et al., 1997; Gray and Bradley, 2005; King and Sinclair, 2015).

In mosquitoes, the role of eggshell in ERD is related with the reduction in the rate of water loss. The outermost mosquito eggshell layer is the exochorion, a delicate layer that easily detaches from the endochorion and does not participate in ERD (Monnerat et al., 1999; Farnesi et al., 2015). Although the endochorion visibly melanizes, the serosal cuticle below it might also do so. In previous works our group have shown images of transparent serosal cuticles from different mosquito species (Rezende et al., 2008; Goltsev et al., 2009; Vargas et al., 2014; Farnesi et al., 2015). However, these cuticles were obtained through bleach treatment, that digests the chorion. During this process, the bleach-resistant serosal cuticle might get unpigmented. In the mosquito *An. gambiae*, the serosal cells, which produce the serosal cuticle, express *tyrosine hydroxylase* and *dopa decarboxilase* genes (Goltsev et al., 2009), coding for enzymes related with both melanization and sclerotization pathways (Figure 1B) (Wittkopp and Beldade, 2009; Arakane et al., 2016). Beckel demonstrates that mosquito eggs without exo and endochorion exhibit a permeable serosal cuticle. Together with the known permeability of eggs before secretion of the serosal cuticle, it seems that the endochorion-serosal cuticle bonding is the functional entity responsible for reducing water loss (Beckel, 1958). This bounding would occur through crosslinking quinones derived from the sclerotization or through interactions with melanins (Andersen, 2012; Arakane et al., 2016).

The moderate level of ERD before serosal cuticle formation, shown in *Ae. aegypti* and *Cx. quinquefasciatus*, but not in *Anopheles* spp., cannot be related to the presence of melanin. This viability might be due to an increased tolerance to water loss or a higher initial egg water content. Percentage of eggshell weight in relation to total egg weight indeed suggest that total body water content is lower in *An. aquasalis* (Farnesi et al., 2015).

Notwithstanding, color traits related with the decrease in water loss evolve differentially in other insect eggs. The eggshells of the cricket *Acheta domesticus* and the beetle *Tribolium castaneum* are transparent. In *A. domesticus* the molecules dopa, dopamine and NADA, (Figure 1B), are present in the serosal cells and cuticle most likely participating in the sclerotization pathway (Furneaux and McFarlane, 1965). In *T. castaneum* the serosal cuticle is fundamental for ERD (Jacobs et al., 2013) and gene silencing of *Laccase2*, related with both melanization and sclerotization (Figure 1B) (Arakane et al., 2016) diminishes the ERD level of this beetle (Jacobs et al., 2015).

### Evolution and ecology of resistance to desiccation in mosquito eggs

Mosquitoes of *Aedes*, *Culex* and *Anopheles* genus shared a last common ancestor ~217 million years ago. The subfamilies Culicinae (containing *Aedes* and *Culex* genera) and Anophelinae have separated ~204 million years ago (Reidenbach et al., 2009). Within this time span the level of pigmentation has greatly diverged, to the point where *Ae. aegypti* and *Cx. quinquefasciatus*, more closely related than *Anopheles* species, show the highest divergence in levels of egg pigmentation and desiccation resistance.

In mosquitoes, egg resistance to desiccation is a trait that guarantees survival in hostile environments and enables population growth and spread to new habitats (Juliano and Lounibos, 2005; Brown et al., 2011). In the case of *Ae. aegypti*, with a high ERD, this implicates in vector dispersion and promotes transmission of diseases such as chikungunya (Vega-Rua et al., 2014), dengue (Bhatt et al., 2013) and Zika (Freitas et al., 2016). Mosquito species with increased ERD are contained in a few genera (*Aedes*, *Haemagogus*, *Ochlerotatus*, *Opifex* and *Psorophora*), adding to about 30% of all described species (Juliano and Lounibos, 2005).

*Aedes aegypti* shows an outstanding success in keeping its eggs viable outside the water, up to 8 months in the dry (Christophers, 1960; Clements, 1992). There is even a report that shows hatching of *Ae. aegypti* eggs after 15 months, when kept at 9 ºC (Bacot, 1918). Indeed, more detailed analysis reveals this hatching success is directly related to higher relative humidity (Kliewer, 1961). The present results show that the increased *Ae. aegypti* eggshell melanization is one of the traits responsible for the extremely efficient ERD seen in this species (Figure 6).

**Figure 6:**
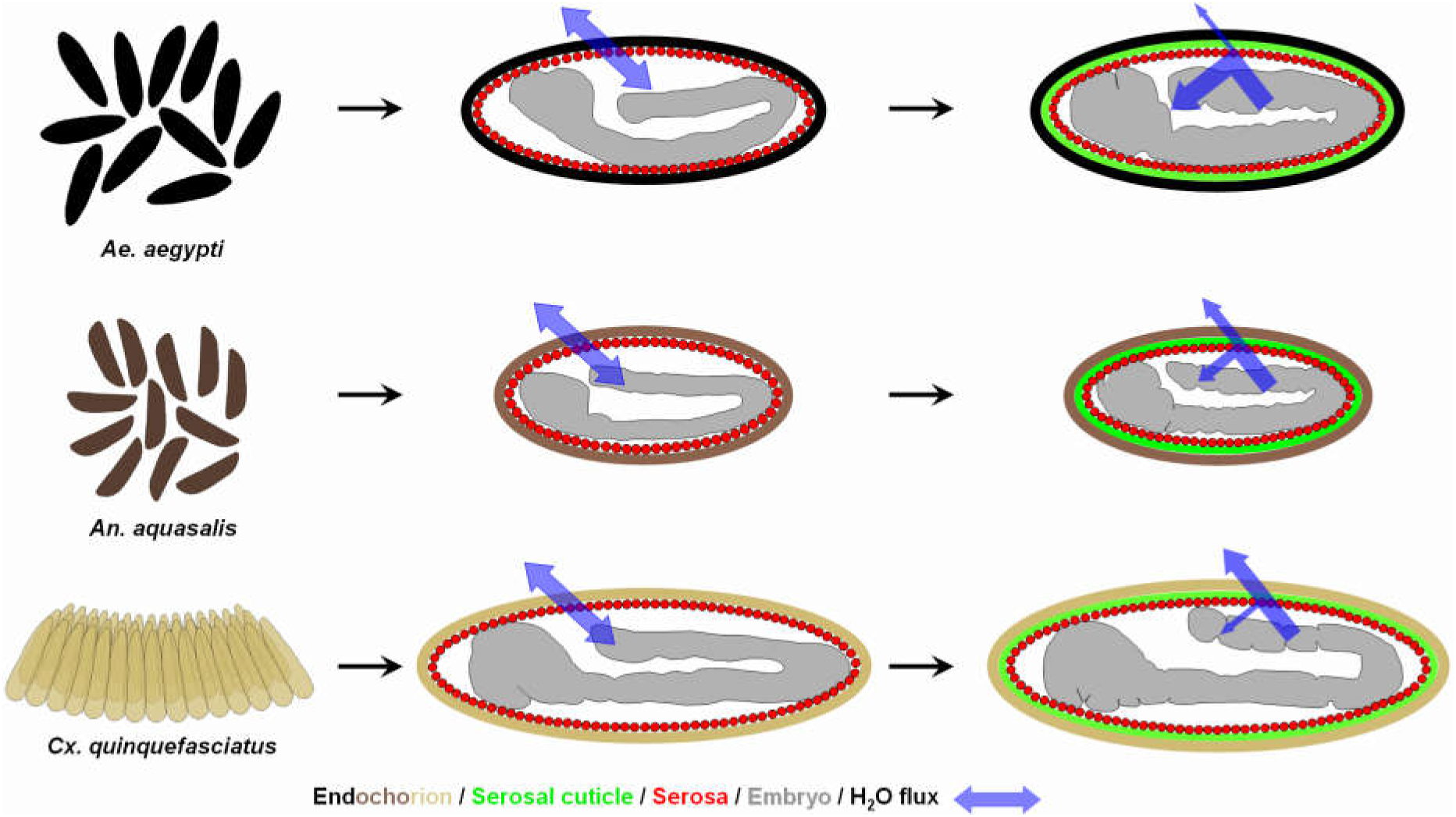
Mosquito vectors egglaying behavior and water flux through the eggshell before and after serosal cuticle formation. From top to bottom, leftmost panel: while *Ae. aegypti* and *An. aquasalis* females lay their eggs individually, the females of *Cx. quinquefasciatus* lay their eggs as an organized raft that floats on the water surface. In all species, before serosal cuticle formation water passes freely through the eggshell. Serosal cuticle formation diminished water passage through the eggshell in a color-dependent manner: while in *Ae. aegypti*, with a black endochorion, most of the water is retained inside the egg, in *An. aquasalis,* with a dark-brown endochorion, some of the water is retained inside the egg, but not all. Finally, in *Cx. quinquefasciatus*, with a light-brown/light-tanned endochorion, most of the water escapes and only a small portion of it is retained inside the egg. The depicted embryonic morphology are representative for each stage and species (Vargas et al., 2014) and egg sizes among species are depicted in their natural proportion (Farnesi et al., 2015). For the sake of simplicity, the outermost eggshell layer (the endochorion) and the other extraembryonic membrane (the amnion) are not depicted here. The exochorion does not participate in the ERD (Farnesi et al., 2015).

Although species from other genera such as *Culex* and *Anopheles* show a less striking ERD (Clements, 1992; Vargas et al., 2014), this trait might still be relevant for survival, at least for Anopheline species. Eggs of *Anopheles* mosquitoes are viable on a dry surface for approximately one day after the end of embryogenesis (Figures 1C and 6) (Darrow, 1949; Vargas et al., 2014). However, when left at humid soil, egg viability increases up to 7 and 18 days in *An. quadrimaculatus* and *An. arabiensis*, respectively (Deane and Causey, 1943; Kartman and Repass, 1952; Parmakelis et al., 2008); other species resist for even longer periods (Clements, 1992). Anopheline egg survival in soil is crucial for sustaining the mosquito life cycle during the dry season and thus the maintenance of malaria outbreaks (Stone and Reynolds, 1939; Beier et al., 1990; Shililu et al., 2004). Moreover, adults from species of the *Anopheles gambiae* complex show distinct levels of resistance to desiccation (Gray and Bradley, 2005; Lehmann et al., 2010). As a future prospect, it would be interesting to evaluate if these species have distinct levels of melanization in their eggshells and adult cuticles.

Females of *Culex quinquefasciatus* oviposit in rafts containing from few dozens to hundreds of eggs arranged along their longitudinal axis. Eggs internal to the raft structure bear sides protected by contact with other eggs; their anterior region contacts the water film, and the posterior tip is the only region in contact with the air (Christophers, 1945; Clements, 1992). Beyond being darker than other eggshell regions (Figure 2), the posterior tip is the only endochorion region whose surface is rough and irregular, similar to the whole endochorion of *Ae. aegypti* eggshells (Farnesi et al., 2015). Given that Culex eggs at raft edges were found dead after exposure to strong dry winds (Clements, 1992), it seems that the raft per se can act as a protection against dehydration, according to the egg cluster-desiccation hypothesis (Clark and Faeth, 1998). This could relax the selection pressure of other traits related with EDR, such as serosal cuticle efficiency and eggshell pigmentation (Figure 6), with the exception of the posterior tip. The occurrence of a higher rate of water loss through the *Culex* eggshell might be advantageous, in the context of a more efficient gas exchange and a increased defense against pathogens, as previously discussed (Vargas et al., 2014).

In summary, we believe that the differential egg resistance to desiccation observed in distinct mosquito species is a trait with multifactorial origins. Eggshell melanization and serosal cuticle formation increases this protection (Figure 6). However, other factors might also contribute such as the thickness and texture of the distinct eggshell layers and the parental investment, observed in *Culex* species.

## Conclusions

Our results demonstrate that, in mosquitoes, the eggshell melanization level is directly associated with egg viability outside the water after serosal cuticle formation. Decoding the association between egg coloration and resistance to desiccation is relevant for studies concerning ecology and evolution of mosquitoes and other insects. Since eggshell and adult cuticle pigmentation ensure survivorship for some insects, they should be considered regarding species fitness and also for the control and management of vector or pest insects (Semensi and Sugumaran, 1986; Prasain et al., 2012).

## List of symbols and abbreviations

DHI: 5,6-dihydroxyindole
DHICA: 5,6-dihydroxyindole-2-carboxylic acid
ERD: Egg resistance to desiccation
GORO: GOlden cuticle + ROse eyes
α-MDH: (DL)-3-(3,4-dihydroxyphenyl)-2-hydrazino-2-methylpropionic acid
NADA: N-acetyldopamine
NBAD: N-β-alanyldopamine

## Acknowledgments

HCMV and LCF were both fellows supported by CNPq grants. The following mosquito strains were obtained through the MR4 as part of the BEI Resources Repository, NIAID, NIH: *Anopheles quadrimaculatus* ORLANDO, MRA-139 and *Anopheles quadrimaculatus* GORO, MRA-891, both deposited by MQ Benedict. We thank Dr. Phil Lounibos and the staff of the FMEL for the space and all assistance for the accomplishment of the experiments with *An. quadrimaculatus*. We thank Maria Cristina Carrasquilha, Tanise Stenn, Erick Blosser and Gabriela Maxxine for the assistance in rearing the *An. quadrimaculatus* strains. We thank all the staff at LAFICAVE for the assistance in obtaining eggs and adults of *Ae. aegypti*, *An. aquasalis* and *Cx. quinquefasciatus* and Luciana Araripe for critical reading and suggestions on the manuscript.

### Competing interests

The authors declare that they have no competing interests

### Author contributions

Conceptualization, DV, GLR; Methodology, DV, GLR; Validation, LCF, GLR; Formal analysis, LCF, HCMV; Investigation, LCF, HCMV; Resources, DV, GLR; Writing - original draft, LCF, HCMV, DV, GLR; Writing - review & editing, DV, GLR; Visualization, GLR; Supervision, DV, GLR; Project administration, DV, GLR; Funding acquisition DV, GLR.

### Funding

This work was supported by FAPERJ (E-26/170.025/2008 and E-26/101.531/2010 to DV, E-26/111.978/2012 and E 26/111.238/2014 to GLR) and CNPq (573959/2008-0 to DV). LCF and HCMV were fellows from CNPq.

